# Establishing an in vitro pipeline for the high-throughput quantification of epithelial permeability of gut bacterial metabolites

**DOI:** 10.1101/2025.06.10.658935

**Authors:** Amber Brauer-Nikonow, Carlos Geert Pieter Voogdt, Nathan Vinícius Ribeiro, Michael Zimmermann

## Abstract

The epithelium of the human gastrointestinal tract is key for controlling the absorption of small molecules and for forming a tight barrier between the gut microbiota and the host, thereby maintaining metabolic homeostasis. Both the microbiota and the barrier function of the intestinal epithelium have a role in pharmacokinetic variability of medical drugs. In this study, we developed a high-throughput workflow to assess the absorption of bacterially produced (drug) metabolites by intestinal epithelial cells through the combination of anaerobic bacterial cultures and human-intestine derived Caco2 cell cultures in a transwell system. To functionally monitor the barrier integrity during the experiments, we introduced a panel of marker compounds, whose concentration kinetics on either side of the epithelial monolayer indicates barrier integrity and transport. We employed this workflow to systematically probe the effect of different gut bacterial species on the epithelial absorption of 482 drugs and their 172 bacterially produced metabolites. While we could recapitulate known bacterial drug biotransformation reactions and expected drug metabolite absorption profiles, we also identified 33 new bacteria-drug pairs for which bacterial biotransformation alters epithelial permeability. Further, we combined the developed experimental workflow with untargeted metabolomics analysis to systematically study epithelial permeability of metabolites naturally produced by gut bacteria. Tracking the absorption kinetics of 397 bacterially produced metabolites revealed that the majority (>79%) of these metabolites do not pass the Caco2 monolayer, illustrating its role as a physical and metabolic barrier. In summary, we present a highly adaptable high-throughput workflow to quantitatively study the metabolic interactions at the intestinal microbiota-host interface which can impact pharmacokinetics, toxicokinetics, and human physiology.

## Introduction

The intestine is the second largest extended surface between the environment and the human body (Gallo, 2017). The complex structure of the intestinal barrier regulates which compounds from the luminal contents enter systemic circulation (Haroun et al., 2023). Moreover, the intestinal barrier is involved in maintaining metabolic homeostasis, which is influenced by the composition and activity of the gut microbiota, the immune system, hormonal state, and activity of intercellular connections of the host (Di Vincenzo et al., 2024). The epithelial layer of the intestinal barrier controls the absorption of water and hydrophilic substances through tight junction proteins (Chelakkot et al., 2018). In contrast, small hydrophobic compounds can be absorbed by the transcellular route, which involves passive diffusion through the cells (Schoultz & Keita, 2020). Many orally administered drugs are absorbed through this latter route, which has also been described by the empirical Lipinski rule of 5 (Lipinski, 2000). Certain compounds are transported actively by the intestinal epithelial cells, for example, water-soluble vitamins, ions, amino acids, and short chain fatty acids. This active transport is mediated by carrier and transport proteins (Schwenk, 1987). For orally administered drugs to be systemically active, the compound or its metabolites have to pass through the intestinal barrier.

The large number of different microorganisms colonizing the human gastrointestinal tract (GIT), also called the gut microbiota, play a significant role in the determination of host physiology (Fan & Pedersen, 2021). Moreover, the gut microbiota can directly metabolise drugs affecting their pharmacokinetics (Verdegaal & Goodman, 2024; Weersma et al., 2020). Through interactions of bacteria with drug molecules in the intestine, the compounds can be biotransformed into different drug metabolites (Zimmermann et al., 2019a), which may have a similar or altered activity compared to the parent drug. For example, the prodrug sulfasalazine is activated by azoreductases, whereas brivudine can be biotransformed into the liver toxic compound bromovinyl uracil, which are both reactions that can be performed by both gut bacteria and human cells (Peppercorn & Goldman, 1972; Zimmermann et al., 2019b). The interpersonal differences in the gut microbiota composition and metabolic activity have therefore been suggested to potentially affect drug pharmacokinetics and pharmacodynamics (Javdan et al., 2020; Zimmermann et al., 2019a). Several anecdotal cases of gut microbial drug biotransformation that impact drug disposition have been described. For example, for chloramphenicol, levodopa, and methylprednisolone (Arassi et al., 2024; Glazko et al., 1952; Isildar et al., 1988; Maini Rekdal et al., 2019). However, to systematically assess the effects of the gut microbiota on systemic drug and metabolite exposure, we currently lack high-throughput approaches to quantify gut microbial contributions to specific processes related to drug absorption, distribution, metabolism, excretion, and toxicity (ADMET).

To quantify the apparent permeability of bacterial metabolites, drugs, and bacterially produced drug metabolites across the colonic epithelial barrier as a factor of oral bioavailabililty, we combined bacterial cultivation with a Caco2 cell monolayer cultures in a transwell system and with high-throughput liquid chromatography coupled to mass spectrometry (LC-MS) analysis. We measured the transfer across the colonic barrier of 482 drugs, 172 gut bacteria-produced drug metabolites, and identified 33 bacterial drug metabolites with altered absorption properties compared to their respective parent drugs. Beyond the scope of medical drugs, we furthermore, employed an untargeted metabolomics approach to monitor the kinetics of 7240 bacterial metabolites, and demonstrate the generalizability of the developed protocol to quantify metabolic microbiota-host interaction. Altogether, we established a high-throughput workflow to mechanistically investigate gut microbial impact on intestinal metabolite absorption that can lead to altered metabolic microbiota-host interactions and interpersonal variability in drug pharmacokinetics.

## Results

### Establishment of a colonic transwell system that includes bacterial metabolites

To quantify the permeability of gut bacteria-produced metabolites, we combined anaerobic cultivation of human gut bacteria with a cell culture-based transwell assay (Figure 1A). We used Caco2 cells, which are a heterogenous population of human intestinal epithelial cells that form a closed cell layer barrier upon cultivation on a transwell membrane. The use of these cells has been well-established for permeability assays of drugs and other compounds. They have become a standard in preclinical drug discovery and development, as evidenced by more than 4400 permeability assays using Caco2 cells currently reported in the ChEMBL data repository (Lennernäs et al., 1996; van Breemen & Li, 2005). To test the effect of gut bacteria on intestinal permeability, we anaerobically cultivated ten different gut bacterial species covering three major phyla (Actinomycetota, Bacteroidota, Bacillota) of the human gastrointestinal microbiome. Importantly, the selected bacterial species have previously been shown to biotransform a broad range of chemicals, such as drugs and carcinogens (B. Zhang et al., 2024; Zimmermann et al., 2019a).

**Figure 1:**
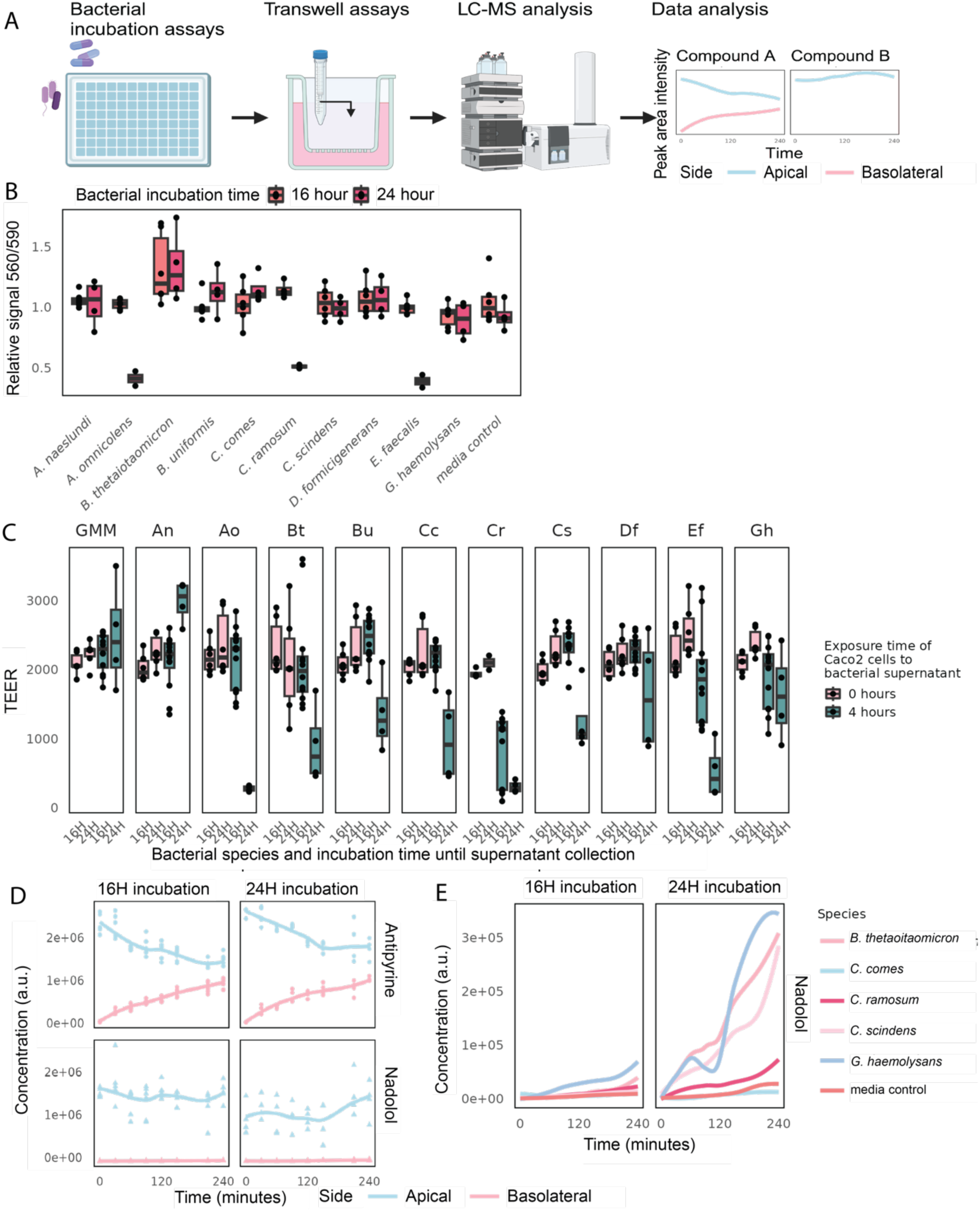
Workflow for measuring bacterial metabolites across an intestinal barrier. A. Schematic overview of the workflow. Created in Biorender.com. B. Viability measurements using resazurin reduction of Caco2 cells on a transwell inserts after 4-hour exposure to bacterial culture supernatants (n=4). Values are depicted as relative changes to the control (sterile bacterial culture media, GMM) performed for each independent experiment. C. TEER values of Caco2 cell monolayers on transwell inserts before and after 4 hours of incubation with bacterial culture supernatants collected after either 16 or 24 hours of bacterial growth (n=2-6). D. Apical and basolateral kinetics of two example marker compounds, antipyrine (permeable) and nadolol (non-permeable), measured upon exposure of Caco2 cells to sterile GMM over 4 hours (n=4). E. Basolateral kinetics of nadolol, a minimally permeable marker indicating defective barrier integrity after exposure of Caco2 cells to bacterial culture supernatants after 24-hour of bacterial growth (n=4). Mean values of the replicates are shown. Abbreviations: GMM: sterile media control, An: *A. naeslundi*, Ao: *A. omnicolens*, Bt: *B. thetaiotaomicron*, Bu: *B. uniformis*, Cc: *C. comes*, Cr: *C. ramosum*, Cs: *C. scindens*, Df: *D. formicigenerans*, Ef: *E. faecalis*, Gh: *G. haemolysans*.

To test whether these bacteria affect Caco2 viability and barrier integrity, we cultivated the bacteria under anaerobic conditions in gut microbiota medium (GMM) (Goodman et al., 2011) for 16 and 24 hours. We then filter-sterilised (0.22 µm) the bacterial culture supernatants and added these supernatants containing bacterial metabolites to the upper compartment of a transwell insert covered with a Caco2 cell monolayer. In this experimental setup, the upper compartment of the transwell is seen as the apical side of the intestinal barrier, which faces the intestinal lumen and is hence directly exposed to gut bacterial metabolites. Correspondingly, the lower compartment of the transwell setup represents the basolateral side of the intestinal barrier, facing the intestinal tissue (Figure 1A). Caco2 cells were exposed to sterile-filtered bacterial culture supernatant (CFS) for four hours and we assessed Caco2 viability by resazurin reduction (indicating metabolic activity by the cells) and barrier integrity by transepithelial electrical resistance (TEER) measurements. Caco2 cell viability was indifferent following exposure to most bacterial culture supernatants cultivated for 16 hours compared to the media control. However, CFS of some species grown longer (24 hours) did impact cell viability (Figure 1B). TEER is a standard measurement of barrier establishment, in which higher values indicate a tighter epithelial barrier. However, its usefulness to assess the monolayer integrity has been debated in literature (Mukherjee et al., 2004; Narai et al., 1997). We found that exposure to specific bacterial culture supernatants (*e.g.*, *Alloscardovia omnicolens*, *Clostridium ramosum*, and *Enterococcus faecalis*) decreased the TEER (Figure 1C). These results made us wonder how bacterial metabolites affect the permeability of the barrier for small molecules.

To directly assess permeability through the intestinal cell layer, we assembled a mixture of seven marker compounds that we added to the apical side of the Caco2 monolayer following their pre-conditioning with CFS of each of the 10 bacterial species. We selected four marker compounds that readily pass through the intestinal barrier by passive diffusion (antipyrine (Lennernäs et al., 1996), haloperidol (Niemegeers & Laduron, 1976), propranolol (Wang et al., 2010), and warfarin (O’Reilly et al., 1963)), and three compounds that poorly pass through an intact epithelial barrier to the basolateral compartment of the transwell system (etoposide (Leu & Huang, 1995), nadolol (Sasaki et al., 2022), and terbutaline (Lennernäs et al., 1996)). We then measured the levels of the marker compounds on either side of the monolayer every 30 minutes for four hours using LC-MS. The resulting marker compound kinetics in the sterile media controls demonstrate the distinct absorption properties of the selected marker compounds (Figure 1D, Figure S1). Importantly, low or absent signal for non-permeable compounds on the basolateral side demonstrated that the function of the barrier remained intact after exposure to supernatants of nine and seven species at 16 hours and 24 hours of bacterial growth, respectively (Figure 1E and Figure S2A). Moreover, using these marker compounds, we found that barrier function does not necessarily correlate with altered TEER values (*e.g.*, for *Gemella haemolysans* and *E. faecalis*) (Figure 1B-E).

Of note, the difference in TEER values, cell viability, and compound absorption between the 16- and 24-hour bacterial cultures was striking for some of the bacterial species tested (Figure 1B-C). We found that the culture supernatants of the 24-hour cultures of *A. omnicolens*, *C. ramosum*, and *E. faecalis*, exerted the strongest impact on Caco2 barrier integrity. The low pH values of these supernatants (pH 5.0) compared to pH 6.9 of the other culture supernatants and the media control suggested a potential pH effect on the epithelial barrier. Indeed, re-adjusting the pH of the supernatants of *A. omnicolens*, *C. ramosum*, and *E. faecalis* cultures from 5.0 to 7.0 negated their negative effect on Caco2 cells (Figure S2B). These results demonstrate that a lower pH, as a result of gut bacterial fermentation, can have an effect on Caco2 viability and barrier integrity of the monolayer. As a consequence, we employed bacterial cultures grown for less than 16 hours and monitored the pH of bacterial culture supernatants for all subsequent assays.

Taken together, these experiments demonstrate that the established experimental setup allows the exposure of Caco2 monolayers to bacterial metabolites and the monitoring of the barrier function. Further, we identified two cases, for which the TEER was minimally impacted while the marker compounds revealed an impaired barrier function (*G. haemolysans* and *C. scindens*). Thus, the employed set of the marker compounds provides a direct and sensitive functional readout of the intestinal barrier function, that is more specific and relevant than TEER measurements.

### Bacterial biotransformation assay compatible with human cell cultures

To study the epithelial permeability of gut bacterial drug metabolites, we sought to establish a high-throughput gut bacterial drug biotransformation assay that is compatible with human cell culture. To this aim, we selected 482 chemically diverse compounds with a broad range of predicted compound hydrophobicity (aLogP) (Figure S3A-B, Table S1). To quantify the permeability of these 482 drugs, we incubated Caco2 monolayers in the transwell setup with the drugs in a combinatorial pooling scheme in quadruplicate for four hours (Zhang et al., 2024; Zimmermann et al., 2019a). Measurements of the kinetics of the seven marker compounds for each transwell and resazurin reduction following drug incubation demonstrated that none of the compounds damaged the epithelial barrier or affected Caco2 viability at the applied concentration of 5 μM (Figure 2A, Figure S3C). Quantification of the drug kinetics on both apical and the basolateral side of the epithelial layer revealed that 373 of the 482 tested compounds passed the epithelial layer to the basolateral side (Figure 2B). This is in line with the expected behaviour of the selected drugs, most of which are approved for oral administration.

**Figure 2:**
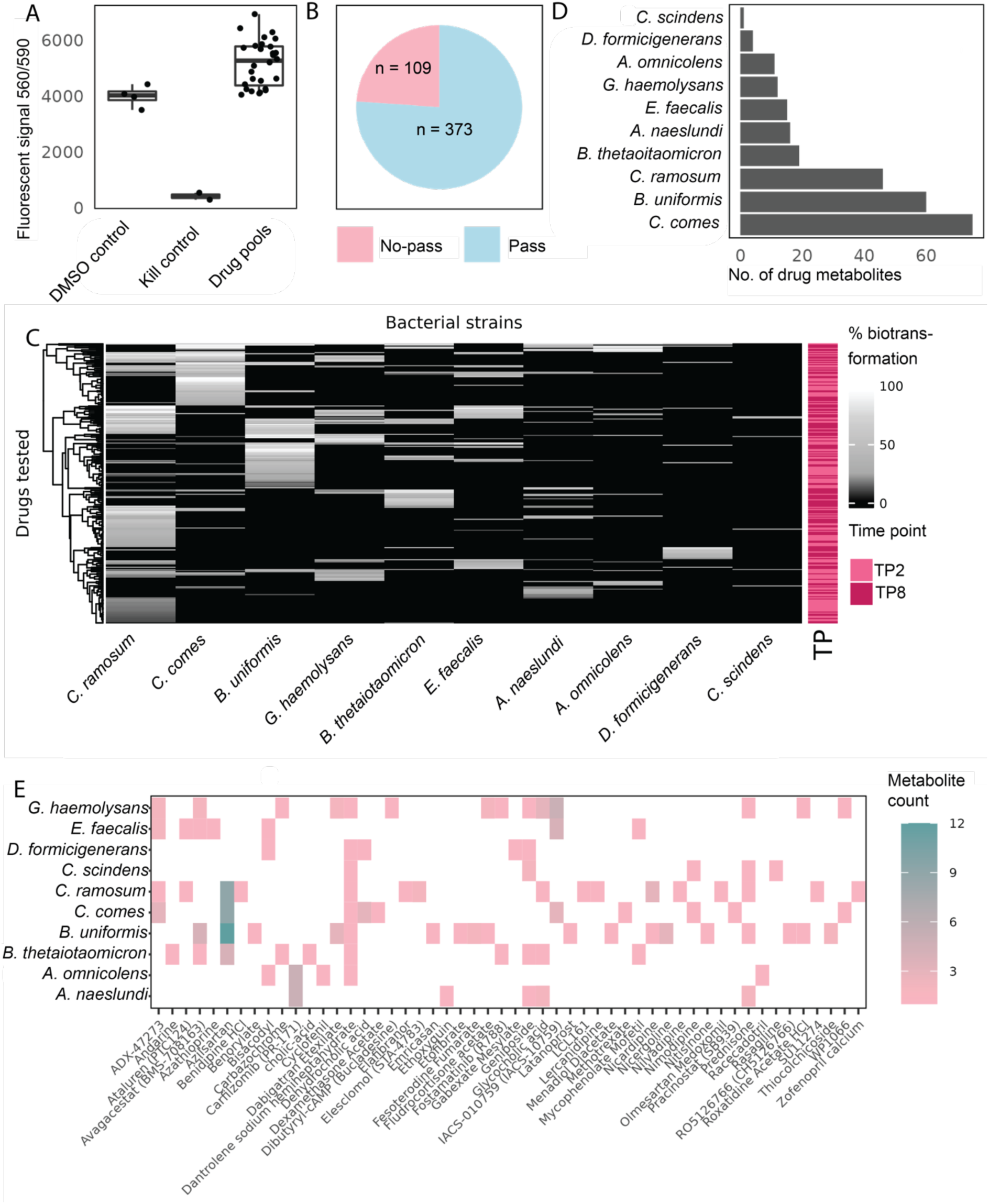
Quantifying epithelial permeability in the transwells and biotransformation of drugs by bacteria of the microbiota. A. Caco2 cell viability after exposure to the drug pools for four hours. B. Drug behaviour in the transwell assay. Drugs were categorised as passing the Caco2 monolayer by subtracting the area under the curve (AUC) from time points 150-240 minutes from the AUC of time points 0-120 minutes (n=4 per drug). C. Heatmap of drug biotransformation by the different gut bacterial strains. The shade of pink described whether a drug was biotransformed after 2 hours, or at 8 hours of incubation. The data from the drugs is clustered based on Euclidian distance (n=4 per species-drug interaction). The species have been sorted based on the number of drugs metabolized. D. Number of putative drug metabolites identified and integrated in the transwell experiment per gut bacterial species tested. E. Heatmap of putative drug metabolites measured in the transwell assays per drug and species combination (n=4 per species-drug).

Next, we incubated these drugs with the ten different gut bacterial cultures described above (and no-bacteria controls) in quadruplicate using the combinatorial pooling protocol previously described (B. Zhang et al., 2024; Zimmermann et al., 2019a). In a total of 1056 samples, we assessed 4820 bacteria-drug interactions by measuring the compound levels at 0, 2, and 8 hours of incubation using LC-MS. We considered compounds being biotransformed when their levels were significantly decreased over time (>10% reduction, FDR-corrected p-value < 0.05) compared to time zero of the incubation and the no-bacteria control. We found 172 (35.6%) of the 482 tested compounds to be metabolised by at least one of the tested bacterial species. As previously reported, we observed large differences between the tested bacterial species and the drugs they biotransformed (Figure 2C) (Arassi et al., 2024; Zimmermann et al., 2019a). Of the 44 drugs included in this study that overlap with a previous report, we found similar biotransformation results for 61.3% of the drugs. The reason that 16 of the 44 drugs that were biotransformed in a previous report but not in the current screen is likely due to the difference in number of gut bacterial species and strains assayed (10 versus 76) (Zimmermann et al., 2019a). This overlap of drugs to be metabolized by gut bacteria validates the altered biotransformation protocol that we used to increase compatibility with cell cultures. To identify bacterial biotransformation products of the 172 metabolized drugs, we employed untargeted metabolomics analysis of the respective samples. To this aim, we first identified putative metabolites that were increased in the presence of a specific drug compared to control samples (log2(fold change) >2 and FDR-corrected p-value <0.01). We then filtered these putative metabolites based on chromatographic retention, mass defects, and ion fragmentation as previously described (Zimmermann et al., 2019a). We further filtered for metabolites that increased in abundance over the incubation time, while the levels of the respective parent drugs significantly decreased in matching cultures. This analysis yielded a total of 259 putative drug metabolites for further testing of their epithelial permeability (Figure 2D). Consistent with previous studies, distinct mass differences between these putative biotransformation products and their respective parent compounds suggested reduction and hydrolysis reactions to occur most frequently (Zimmermann et al., 2019a) (Figure S4A). Moreover, our biotransformation screen could recapitulate known biotransformation reactions, for example the deacetylation of roxatidine acetate by *B. uniformis* and of bisacodyl by several bacterial species (Javdan et al., 2020; Zimmermann et al., 2019a). We also identified new reactions, such as the N-acylation of azilsartan, an antihypertensive, and the hydrolysis of carfilzomib, an anti-cancer drug that acts as a selective proteasome inhibitor. The latter reaction leads to a drug metabolite that was only described in patient samples previously (Figure S5A-B) (Wang et al., 2013). In summary, employing a high-throughput gut bacterial biotransformation assay with 482 compounds and 10 gut bacterial species allowed us to pinpoint 172 compounds that are biotransformed by at least one of the ten gut bacterial species. Additionally, we identified 259 putative drug biotransformation products for further testing of their permeability through the intestinal epithelial barrier.

### Epithelial permeability of drugs and gut bacterial drug metabolites

Having established the compatibility of the CFS with Caco2 cells and the establishment of the bacterial drug biotransformation assays, we next set out to assess the effect of bacterial (drug) metabolism on trans-epithelial transport. To this aim, we used CFS from the bacterial biotransformation assays at zero and eight hours of drug exposure. The CFS of each drug pool was added to the apical side of a transwell and we collected samples every 30 minutes for four hours from either side of the transwells. In this screen, we analysed 12,672 samples by LC-MS to assess a total of 4,338 drug-bacteria-transport interactions in quadruplicate. We determined the concentration kinetics of each drug and their (putative) bacterial metabolites on either side of the transwell barrier. We then investigated whether any drug metabolite displays different epithelial permeability compared to their parent drug, due to altered physicochemical properties upon gut bacterial biotransformation. We detected 170 of the 259 putative drug metabolites identified in bacterial cultures also in this transwell assay (Figure 2E). To assess the permeability of these compounds across the Caco2 monolayer, we compared the slope of the concentration kinetics on either side of the transwell system for each putative biotransformation product and its originating parent drug. To compare the kinetic profiles, we fit a linear regression model for the apical and basolateral sides for both the parent compound and its putative biotransformation products. We identified 33 (19.4%) pairs of drug and bacterial drug metabolite, for which the permeability kinetics (concentration slopes) were significantly different (FDR-adj. p-val < 0.01) (Table S2). Among these 33 cases, we identified eight cases for which the bacterial metabolite passes the intestinal barrier whereas the parent compound does not and two cases with opposite behaviour. For example, carfilzomib, used in treatment of multiple myeloma, poorly passes the epithelial barrier, while its bacterial metabolite readily passes (Figure 3A). Intriguingly, a previous study reported this metabolite in plasma and urine of patients with solid tumors and multiple myeloma without indicating its potential gut microbial origin (Wang et al., 2013). Further, carfilzomib has a high patient-to-patient variability in its pharmacokinetics, which could hence be (partially) explained by the biotransformation of the drug by the gut microbiota (Brown et al., 2017; Quach et al., 2017). Additionally, we identified nine species-drug-metabolite interactions for which both the bacteria and the Caco2 cells produce the same metabolite. In our screen, one of the drugs that is known to be metabolized by the gut microbiota, bisacodyl, a stimulant laxative, was included. Bisacodyl has to be converted into its active metabolite bis-(p-hydroxyphenyl)-pyridyl-2-methane (BHPM) in the gut (Corsetti et al., 2021) to be active. We found that most bacteria, as well as the Caco2 cells, can convert bisacodyl to BHPM, which readily passes to the basolateral compartment (Figure 3B). Another example for which both bacteria and Caco2 cells biotransform the same drug is nicergoline, which is deacetylated (Figure 3C, Figure S5C). Caco2 cells are active in metabolism and express certain drug-metabolising enzymes (Vaessen et al., 2017), which likely explains the observed biotransformation of some (pro)drugs in the absence of bacterial incubation. In summary, our screen showed that most bacterial drug metabolites have similar epithelial permeability compared to their parent drugs. One of the examples of a drug and metabolite with unchanged absorption properties is the drug azilsartan, a reaction by the bacterial species described above (Figure 3D). Interestingly, we found several bacterial biotransformation products that display altered permeability compared to their parent drugs, which indicates that our workflow can be used to assess the effect of bacterial drug metabolism on altered intestinal permeability of pharmaceuticals. Some of the examples of differential epithelial permeability between bacterial drug metabolites and drugs may contribute to pharmacokinetic variability between patients.

**Figure 3:**
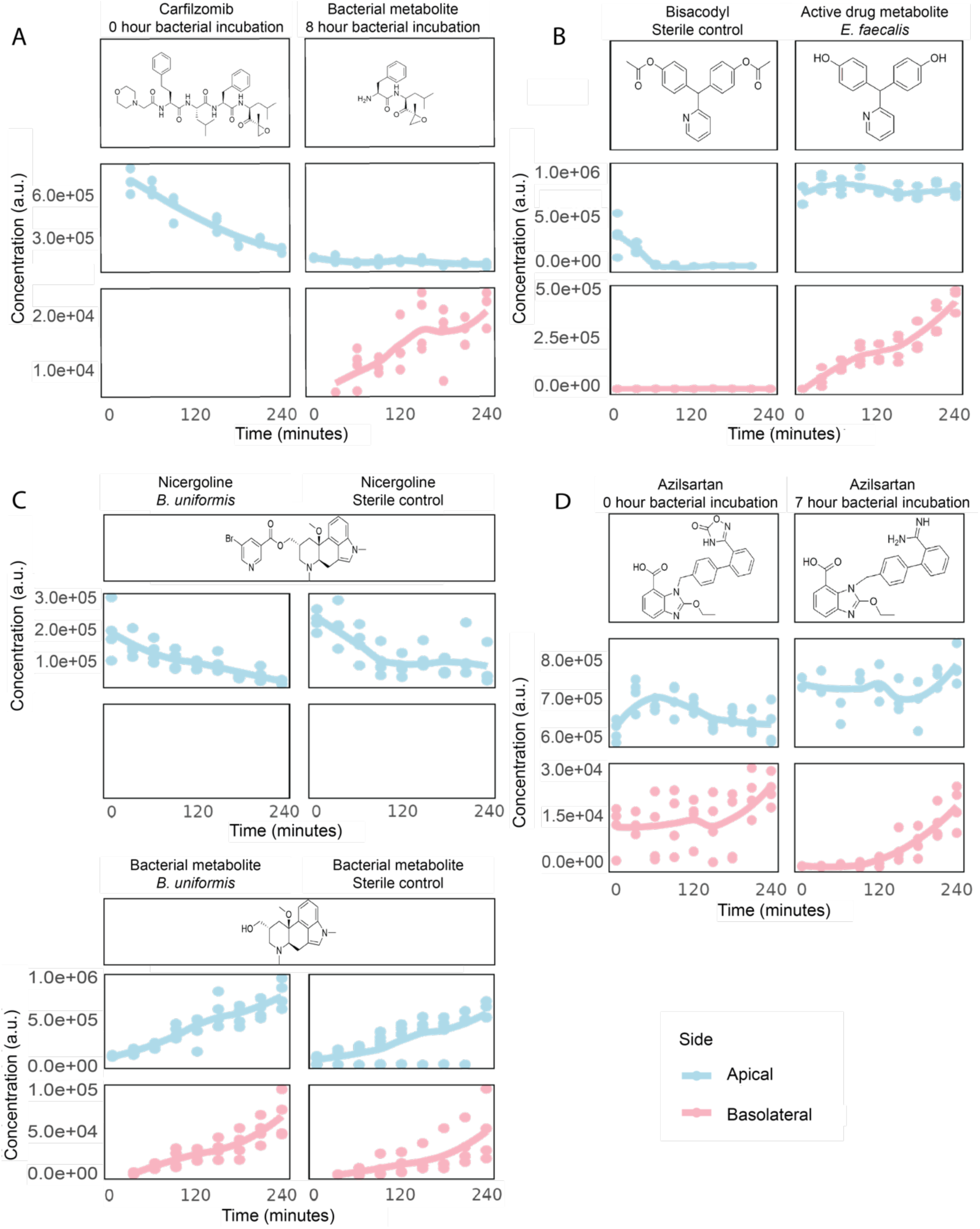
Different cases of drug and drug metabolite absorption behaviours. A. Absorption profiles of carfilzomib and its metabolite, an example a drug in which that does not pass the Caco2 monolayer but the bacterial metabolite can be detected passing to the basolateral side. B-C. Examples of convergent metabolite production by the bacteria and the Caco2 cells. D. Absorption profile of Azilsartan and its hydroxylated bacterial metabolite in the transwell set-up. N=4 replicates per compound for all panels.

### Quantification of intestinal absorption of bacterial metabolites

Next, we sought to employ our established workflow to assess intestinal absorption of endogenous gut bacterial metabolites. To this aim, we employed an untargeted metabolomics approach to identify bacterially produced metabolites and to trace them in the transwell system to assess permeability through the epithelial monolayer. We used the no-drug control data of the screen described above, in which we exposed the Caco2 cells to the supernatant of each gut bacterial species. We annotated (level 3) a total of 7240 out of 13,874 detected metabolic features using the MiMeDB database (Wishart et al., 2023). From these metabolites, we focused on the 10% most abundant metabolites per species that were not detected in the growth media controls, resulting in a total of 1425 metabolites. After fitting these metabolites in the kinetics data from the apical and the basolateral side of the transwell assay, we performed additional filtering steps based on the number of replicates and time points each metabolite was detected in (see materials and methods) (Figure 4A, Table S3). This resulted in a total of 397 bacterially produced metabolites that we detected on the apical side. For 232 of these metabolites we detected no change in their abundance during the four hours of Caco2 cell exposure (Figure 4B). For 104 metabolites, we detected a decrease over time on the apical side (*i.e.,* a negative fold change between time points 0 and 240 minutes). For example, cyclo-(D-arginine-L-proline) produced by *A. omnicolens*, *E. faecalis* and *G. haemolysans* (Figure 4C). Such cyclic dipeptides are commonly secreted by members of the gut microbiota as signalling molecules (Nishanth Kumar et al., 2014; Ogilvie & Czekster, 2023). Of the metabolites that were decreasing on the apical side, we however only detected 19 on the basolateral side with increasing concentrations, suggesting that the metabolite passed unchanged through the Caco2 monolayer. An example is deoxyadenosine (Figure 4D), an adenosine derivative that can be produced by both the host and the microbiota, and that can disrupt the function of immune cells when present at high levels (Boussamet et al., 2024). It has recently been identified as an oral microbial biomarker in a study comparing multiple sclerosis patients and healthy controls (Wishart et al., 2023). In conclusion, our developed workflow can be used to study the (selective) transfer of conserved and unique microbial metabolites across the intestinal barrier in high-throughput. We found many bacterial metabolites that are further metabolized by the Caco2 cells, which show some of the dynamic interactions between the gut microbiota and the intestinal barrier. Together, these findings indicate that only a small number of bacterial metabolites pass unchanged through the intestinal barrier, supporting the notion that the epithelial layer functions selective metabolic barrier and that there is a dynamic metabolic interaction between the gut epithelium and the bacterial metabolites (Di Vincenzo et al., 2024; Ghosh et al., 2021).

**Figure 4:**
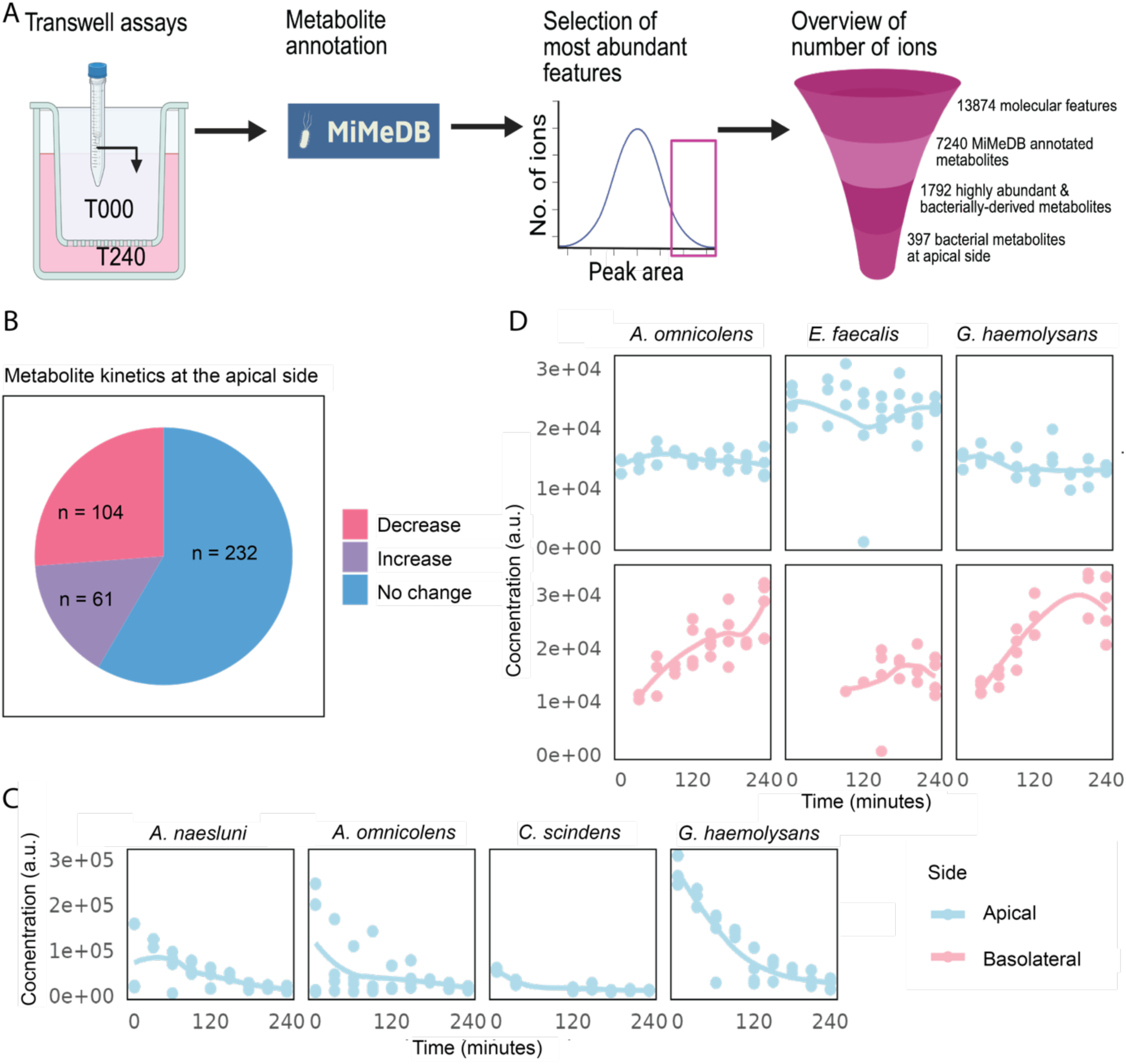
Detecting bacterial metabolites across the epithelial barrier. A. Schematic overview of the data analysis steps to mine the untargeted metabolomics data for bacterially produced metabolites. Created in Biorender.com. B. Pie-chart summarizing the behaviour of the selected bacterial metabolites. The behaviour was determined based on the data at the apical side, for which for each metabolite the log2 fold change of peak area was determined between 0 minutes of exposure and 240 minutes of exposure. A decrease in peak area abundance was defined as a log2fold change value > -0.5, and an increase as a value > 0.5. No change in peak abundance was defined as a log2 fold change value between -0.5 and 0.5 (n=4 per metabolite-species pair). C. Example of cyclo-(D-arginine-L-proline), a metabolite with decreasing levels over the time of incubation on the apical side of the transwell, but that is not crossing to the basolateral side unchanged (n = 4). D. Absorption kinetics of deoxyadenosine, a metabolite, that has been identified to pass the intestinal barrier unchanged (n = 4).

## Discussion

We developed a workflow to study the permeability of microbial (drug) metabolites across the gut epithelial barrier, represented by a Caco2 monolayer on transwell inserts. The established workflow offers a robust, sensitive and high-throughput method to explore how and which bacterial metabolites (selectively) pass the epithelial barrier and how biotransformation may impact pharmacokinetics, and to identify both expected and new drug-microbiota-absorption interactions.

We observed that bacterial metabolites affect TEER values, without impacting Caco2 viability. This indicates that gut barrier integrity is not only a direct function of cell fitness. To monitor subtle changes in barrier integrity, we introduced a panel of marker compounds of epithelial permeability. Applying these marker compounds, we identified cases for which TEER measurements were unchanged, while the barrier function was impaired. It has been previously reported that bacterial metabolites have an effect on the function of tight junction proteins and expression in the epithelial cells through Toll-like receptor signalling (Ey et al., 2009; Ghosh et al., 2021). The differential effects of the supernatants of distinct bacterial species showed the potential role of species-specific metabolites in shaping intestinal barrier function. This could be further explored in the future to gain insights into the molecular effects that the complex bacterial communities of the human gut have on the intestinal barrier functions.

Our high-throughput screen assessing how microbial activity may alter drug absorption revealed several new cases of differential epithelial permeability between drugs and their bacteria-produced metabolites. The identification of bacterial drug metabolites with different absorption properties compared to their respective parent drug suggests a mechanism of how the microbiota can potentially affect pharmacokintetics. However, in our panel, most of the drug-metabolite pairs have comparable absorption properties, while their activity can be different. For example, in the case of the deacetylation of roxatidine acetate (Figure S4B), which forms the active metabolite of the prodrug, both roxatidine acetate and its deacylated form have comparable absorption properties, but the deacetylated metabolite has been described as the active drug (Kang et al., 2021). Moreover, the cases for which the microbiota and the Caco2 cells produced the same metabolite reflects the broader interplay between host and microbiota metabolic pathways and raises the question of potential synergies or multi-step metabolism in drug biotransformation between the host and its microbiota.

The untargeted analysis of Caco2 exposure to endogenous bacterial metabolites highlights the selective nature of the intestinal epithelial barrier to transfer of metabolites, reinforcing its role as metabolic gatekeeper. The untargeted metabolomics approach identified 7240 metabolites, of which 397 were high-abundance, bacteria-derived metabolites. However, only 4.8% (19) of these metabolites could be detected at both the apical and basolateral side, suggesting that most bacterial metabolites do not cross the intestinal barrier unchanged. The decreasing levels of many metabolites on the apical side over time without corresponding detection on the basolateral side suggests their metabolism by Caco2 cells. This is corroborated by our analysis that identified 73 highly abundant basolateral metabolites which were only measured at the basolateral side and that increased over time. This workflow could be used in the future to further investigate the effects of bacterial metabolites on gut barrier function in a high-throughput setup. To mimic the intestinal epithelium better in this system, Caco2 cells could be cultured with air-liquid interface and the addition of vasointestinal peptide (VIP) to enhance apical mucus production (Floor et al., 2025). Moreover, cultures of patient material, intestinal cells from animal disease models, or co-cultures of cells that form a monolayer together, could be used in this experimental system to selectively study the metabolic gut microbiota - gut epithelium interaction. In addition, different cell types could be added to the basolateral compartment of the transwell setup to study the interactions of the gut to other organs, as has been shown before with NCI-H716 enteroendocrine and liver cells (Chaudhari et al., 2021).

In summary, this study highlights the significant role the gut microbiota plays in modulating drug absorption and metabolism, with implications for understanding pharmacokinetic variability among individuals. The developed workflow could be a useful tool in drug discovery and development, as the permeability of drugs across a Caco2 monolayer already is accepted by the FDA as a surrogate for human intestinal permeability measurements. The combination of the permeability studies with the possible interaction of a drug with the gut microbiota, and the effect of absorption of the drug (and drug metabolite) mimics the *in vivo* situation better and can therefore improve drug discovery, development, and regulatory approval processes (Larregieu & Benet, 2013). Therefore, the developed workflow offers a scalable and reproducible method for assessing these interactions, enabling more in-depth studies on the interplay between host cells and microbial metabolites.

### Limitations

This work describes the establishment of a high-throughput workflow to assess absorption properties of microbial (drug) metabolites. In the current setup, some limitations exist. For example, all data in this work has been acquired with a single LC-MS method. Therefore, we likely missed certain (drug) metabolites, since these might not be retained on the selected chromatographic column (*i.e.*, C18). Further, we used Caco2 cells, which are well-established for permeability assays and can mimic the intestinal barrier of the small intestine to some extent, but they are not fully representative of the gut epithelium. The gut epithelium consists of several cell types, which communicate with each other, which has not been included in our experimental setup. Additionally, we used CFS from single species, whereas the gut microbiome consists of a community of different bacterial species. We therefore likely missed metabolic interactions that would happen in the gut. Moreover, the mechanism by which metabolites may change the barrier function of the Caco2 monolayer has not been elucidated in this work. For the bacterial cultures with near physiological pH that nonetheless impaired Caco2 barrier function, it remains unclear how the barrier function exactly is changed.

## Methods

### Chemicals

All chemicals were purchased from SelleckChem. We selected chemicals from the customized chemical library (96-well)-L2000-Z367755-100uL. Combinatorial pools were prepared based on the output of the PoolingScheme package (https://git.embl.de/bartmans/PoolingScheme). The pooling scheme consisted of 28 pools, in which each compound was included in 4 pools (B. Zhang et al., 2024). LC-MS grade solvents were purchased from T.h Geyer and Merck Millipore.

### Bacterial culturing

A flexible anaerobic chamber (Coy Laboratory Products) containing 12% CO2, 2% H2 and 86% N2 was used for all anaerobic microbiology steps. All anaerobic culturing was performed on brain–heart infusion (BHI; Millipore, cat. no 70138) agar supplemented with 10% horse blood (Oxoid Deutschland, cat. no OXSR0050E). Liquid cultures of bacterial gut isolates and whole communities for drug degradation assays were grown in gut microbiota medium (GMM). Cell-free supernatants (CFS) were collected from liquid cultures that were inoculated at OD600 of 0.01 and grown for 16 or 24 hours. The liquid cultures were then filtered sterilized (Millipore, cat. no SLGV033RS) and stored at -80°C until further processing.

### Drug assays

For each drug assay, glycerol stocks of individual bacterial species were plated on a BHI-blood plate and incubated for 24-36 hours. A single colony was selected and inoculated in 10mL pre-reduced GMM. After 24 hours of growth, this culture was diluted and cultured overnight. Upon reaching an OD600 between 0.3 and 0.9, the culture was spun down (5 min, 4337 x g) under exclusion of oxygen and resuspended in pre-reduced diluted GMM (20%) to an OD600 value of 1.0. This culture was incubated with the drugs at a concentration of 4uM until sample collection. At sample collection, each pool was filter sterilised (Agilent filter plate PVDF 0.2uM) and stored at -80°C until further processing for analysis by LC–MS and use in the transwell system.

### Transwell system

Caco-2 cells were obtained from Sigma (86010202) at passage 9 (Caco-2 cells are epithelial cells isolated from a male with colorectal carcinoma). Cells used in this study were not passaged more than 20 times. Transwell inserts were obtained from Millicell, and were used in combination with 24-well plates (Thermo Fisher). The Caco-2 cells were grown in T75 flasks at 37°C in an incubator with an atmosphere of 5% CO2. The cells were grown using Dulbecco’s modified Eagle medium (DMEM) (Gibco, 41965-039) with 4.5 g/liter D-glucose, 584 mg/liter L-glutamine, 1% penicillin/streptomycin, 1% Amphociterin B, and 10% fetal bovine serum. The medium in the flasks was changed every 3 days. At 90% confluence, the cells were passaged using incubation with 0.5% trypsin for 5 minutes at 37°C after washing the cells with PBS. Cells were then seeded into the transwell inserts (pore size 0.4 μm) at 10,000 cells/well for the 96-well plates and at 1e5 cells/well for the 24-well plate inserts (Millicell, PSHT004R1 and PIHP01250). The transwells with the Caco-2 cells were used only after >21 days post-seeding, to allow for the establishment of a stable monolayer. During this post-seeding period, the media was changed every 3 days.

The transwell inserts with the Caco-2 monolayer were washed 2 times with Hank’s Buffered Salt Solution (HBSS), before adding the sterile supernatant from the bacterial strains, or the single compounds in solution to the apical or basolateral side. To ensure the stability of the permeability of the monolayer during the experiment, a panel of 7 drugs at a concentration of 1uM were added together with the samples. Small volumes of both the apical and basolateral side (5 uL) were taken every 30 minutes and immediately stored at -80°C until metabolite extraction. Time points were taken for 4 hours, whereafter the cells were washed three times with HBSS and media was added to the cells, on the apical side containing a viability dye (Promega, G8080). After an overnight incubation, the apical media was transferred to a separate 96-well plate (Thermofisher) and fluorescent signal was measured. Hereafter, cells were given new media for additional assays.

### Mass spectrometry analysis of drugs and metabolites

Samples were prepared for LC–MS analysis by organic solvent extraction as previously described (Zimmermann et al., 2019a). In brief, acetonitrile:methanol (1:1) was added at−20°C after the addition of internal standard mix (sulfamethoxazole, caffeine, ipriflavone, tolfenamic acid each to a final concentration of 80 nM). LC-MS analysis was performed as previously described (Zimmermann et al., 2019a). In brief, chromatographic separation was performed by reversed-phase chromatography (InfinityLab Poroshell HPH-C18, 2,1×100 mm, 1,9 um) using an Agilent 1200 Infinity UHPLC system and mobile phase A (H2O, 0.1% formic acid) and B (methanol, 0.1% formic acid), and the column compartment was kept at 45°C. 5 to 7.5 µL of sample was injected at 100% A and 0.6 mL/min flow, followed by a linear gradient to 95% B over 5.5 min and 0.4 ml/min. The qTOF instrument (Agilent 6546) was operated in positive scanning mode (50–1,000 m/z) with the following source parameters: VCap, 3,500 V; nozzle voltage, 2,000 V; gas temperature, 225 °C; drying gas 13 l/min; nebulizer, 20 psig; sheath gas temperature 225 °C; sheath gas flow 12 l/min. Online mass calibration was performed using a second ionization source and a constant flow (5 μl/min) of reference solution (121.0509 and 922.0098 m/z).

### Data analysis

#### Targeted data analysis drug assays

For targeted analysis of metabolomics data, the data files in the Agilent data format were transformed to .mzML files using Proteowizard (Chambers et al., 2012). To determine the retention time of each of the xenobiotics used in this study, pure compounds were measured and the retention time was defined. These were used in the subsequent experiments in the combinatory pools. For the drug assays, chromatographic peaks of the different drugs were integrated using the R package xcms, with a time window of 20 seconds before and after the previously identified retention time (Tautenhahn et al., 2008). When several peaks of the mass of interest were identified, the peak closest to the originally determined retention time was chosen.

For each bacterial species, drug peak area under the curve (AUC) fold changes were calculated between time points 2 and 0 hours, and 8 and 0 hours in the 4 pools containing the drug. Statistical significance of the fold change was determined using a two-fold t-test (t.test function in r). The reported p-values were adjusted for multiple testing with FDR (p.adjust function, method = “BH”). The same test was performed for the sterile control experiment. Drugs were identified as degraded when the fold change decrease was >10% and the FDR-corrected p-value was <0.1, and if drugs were not identified as degraded in the sterile control.

#### Untargeted data analysis drug assays

For the identification of molecular features as potential drug metabolites, untargeted data analysis was performed. To this aim, the .mzml transformed original data files were analysed with MZmine (Mzmine version 3.7.2) using the batch file in supplementary materials 1 (Heuckeroth et al., 2024). The tables with the molecular features were analysed in R. To this aim, we log-transformed all data and compared the peak AUC for every peak between the pools containing a drug and the pools without by t-test (t.test function in r) at the time point of 8 hours. The reported p-values were adjusted for multiple testing with FDR (p.adjust function, method = “BH”). Molecular features that had an FDR-corrected p-value < 0.01, had a retention time more than 0.4 minutes difference to the parent drug, a m/z value difference greater than 1, were specific to one parent drug only, and were not measured in the sterile control experiment, were selected as potential drug metabolites.

#### Targeted transwell assay data analysis

For the targeted analysis of the transwell data, we also transformed the data files to .mzML format and used xcms to integrate the peaks that were identified earlier. We then selected only the features that met the intensity threshold of 1000 peak intensity area, and that could not be integrated for the sterile control samples. As a result of this data filtering, we integrated 259 putative drug metabolites in the transwell data; filtering for barrier intactness, filtering for at least three technical replicates at at least three different time points, we comprised a dataset containing the absorption profiles of 170 drug-species-metabolite pairs. We then compared the slopes of each parent drug measured from the 0-hour bacterial incubation in the transwell assay and the slope of the drug metabolite measured of the 8-hour bacterial incubation. For 67 of these drug-metabolite pairs, the parent drug had a too low abundance due to degradation by the bacteria, so we used the data from the sterile control experiments to compare the absorption profiles. We used FDR-correction for all p-values collected.

#### Untargeted transwell data analysis

Similar to the drug assay data files, we also used files converted to .mzML, and peak integration was performed using MZmine (Supplementary file 2). Molecular features were annotated to metabolites considering [M+H+] and [M-H-] ions, if no ion mass was calculated by MZmine3, using the metabolite information list from the Kyoto Encyclopedia of Genes and Genomes (KEGG) metabolite repository (Kanehisa et al., 2023). Ions were assigned to metabolites with a mass tolerance of 0.002amu or 20ppm. To identify highly abundant bacterial metabolites, we selected the 10% of the metabolites with the highest mean abundance value across two time points: 0 minutes of exposure at the apical side and 240 minutes exposure at the basolateral side. Quality control filtering was performed with the same parameters as for the targeted analysis.

### Chemoinformatics

Information on the drugs included in the DrugBank database was downloaded from the DrugBank website (go.drugbank.com) on October 11th 2024, which was version 5.1.12., which was released on 2024-03-14. To parse the downloaded xml file, the R package dbparser was used (Ali & Ezzat, 2024). Using this package, the SMILES codes and the calculated LogP values were collected.

Chemical similarity between the drugs from the panel and the compounds included in the DrugBank database (12227 drugs) was calculated based on Morgan fingerprints, by the function AllChem.GetMorganFingerprintAsBitVect function in the RDKit package (Open Source Chemoinformatics, http://www.rdkit.org). These fingerprints were converted into a binary matrix, on which we did a principal component analysis.

### Software, data, and code availability

All data analysis in R was performed using R version 4.2.2., for the peak integrations of the untargeted datasets, MZmine version 3.7.2 was used (batch files are in the supplementary materials). Raw metabolomics data will be made available in the MetaboLights repository, with accession number REQ20250203208391. Code used for analysis of the data can be found on GitLab: https://git.embl.de/grp-zimmermann/transwells_analysis.

## Acknowledgments

We thank Anne-Claude Gavin (University of Geneva) and the members of the Zimmermann group for helpful discussions. This work was supported by the European Molecular Biology Laboratory; the EMBL International PhD Programme (A.BN.); the European Research Council (ERC) (GutTransForm-101078353).

## Author contributions

A.BN. and M.Z. conceived the study. A.BN., C.G.P.V, and N.V.R. conducted experiments. A.BN. analysed and visualized the data. A.BN. and M.Z. wrote the manuscript. All authors approved the final draft of the manuscript.

## Supplementary figures

**Supplementary figure 1:**
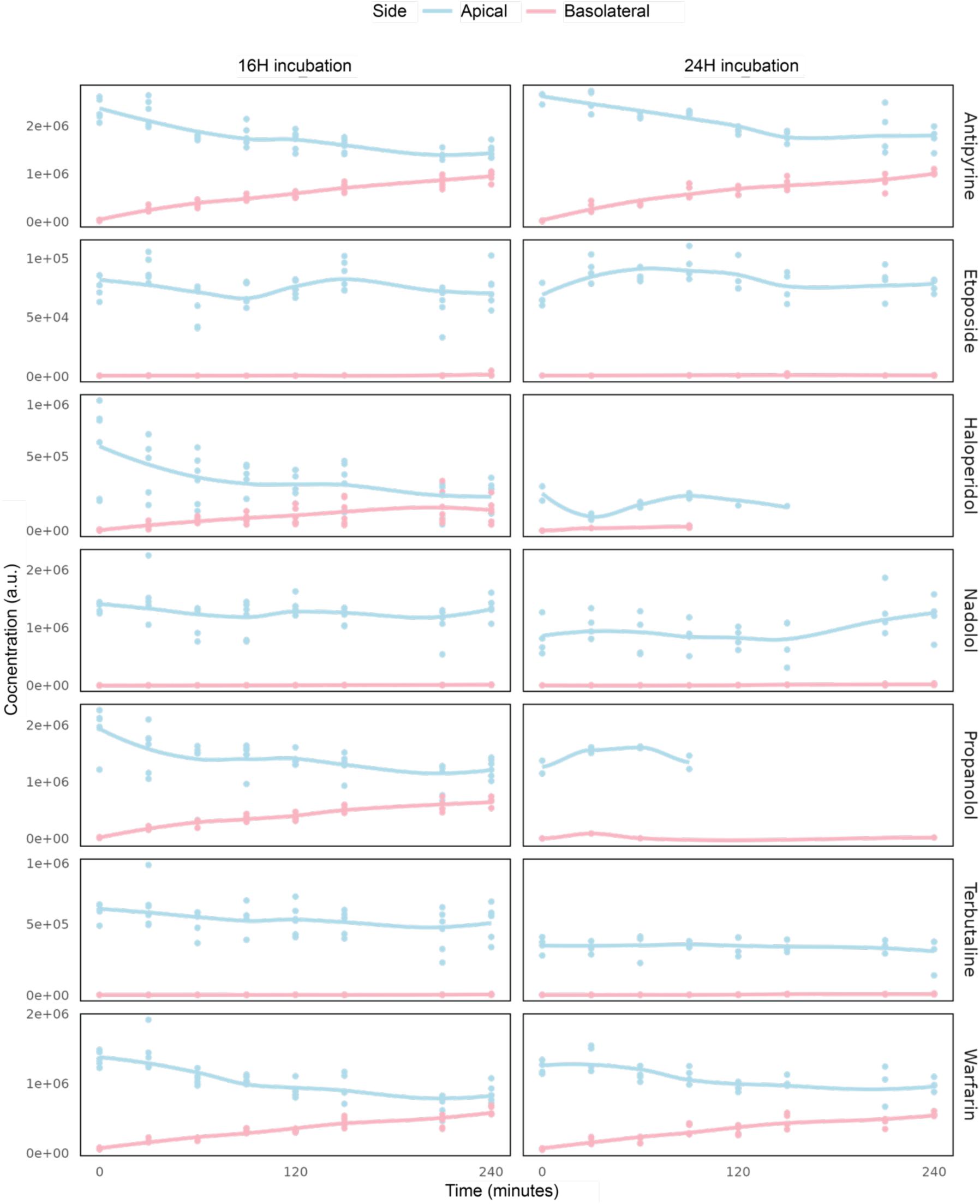
Marker compound kinetics in control conditions (sterile bacterial culture medium) on the apical and basolateral side of the Caco2 barrier. Blue indicates the signal at the apical side, pink indicates the peak areas at the basolateral side (n=4 per time point). Panels on the left are measured with exposure to bacterial supernatants from cultures incubated for 16 hours, and panels on the right with bacterial supernatants from 24 hour incubated cultures.

**Supplementary figure 2:**
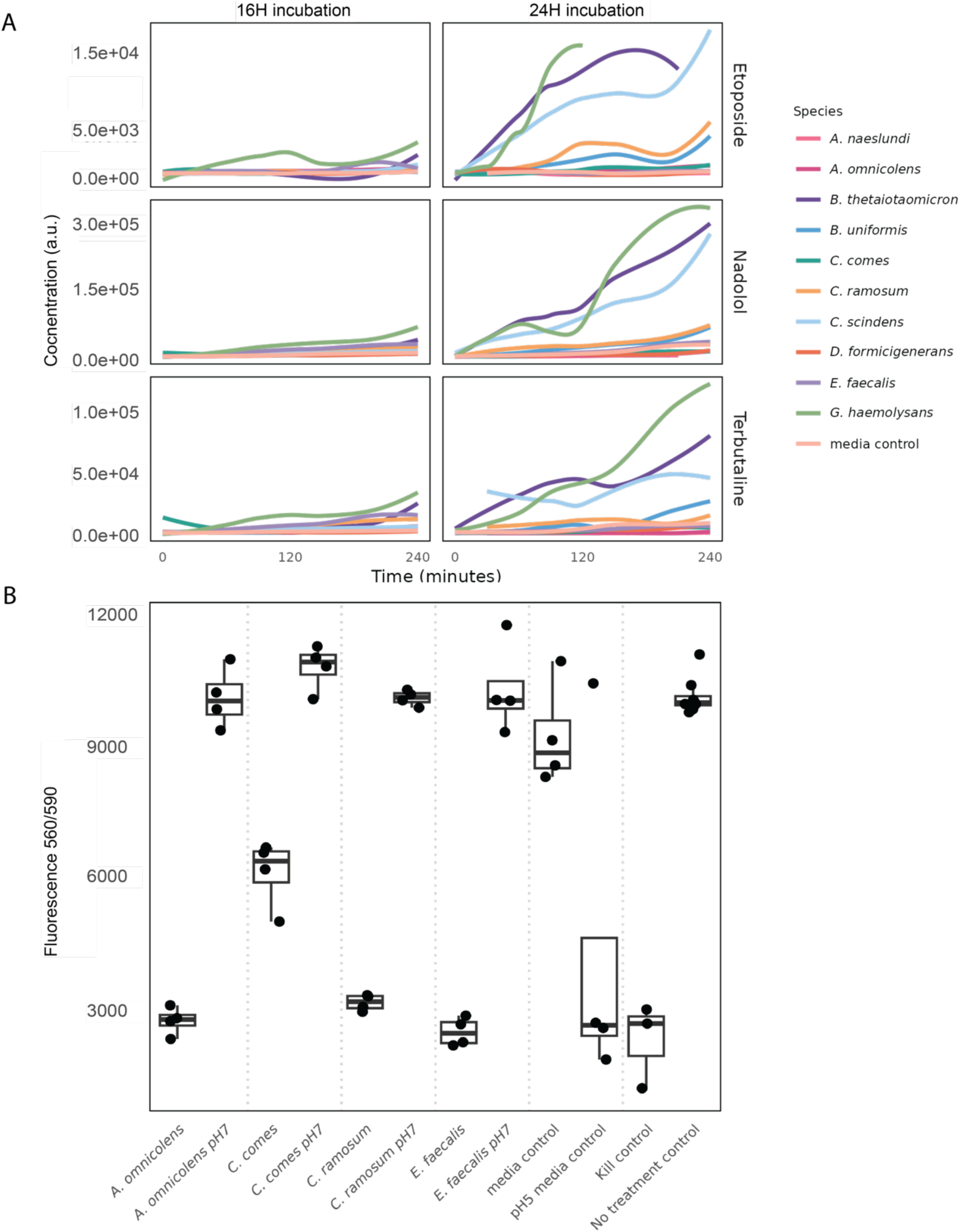
A. Basolateral kinetics of non-permeable marker compounds upon exposure with sterile culture supernatants from 10 different gut bacterial species. Mean values of 4 replicates are shown. B. Caco2 viability after 4 hours of exposure to bacterial culture supernatants after 24 hours of growth before and after pH-adjustment to pH 7.0.

**Supplementary figure 3:**
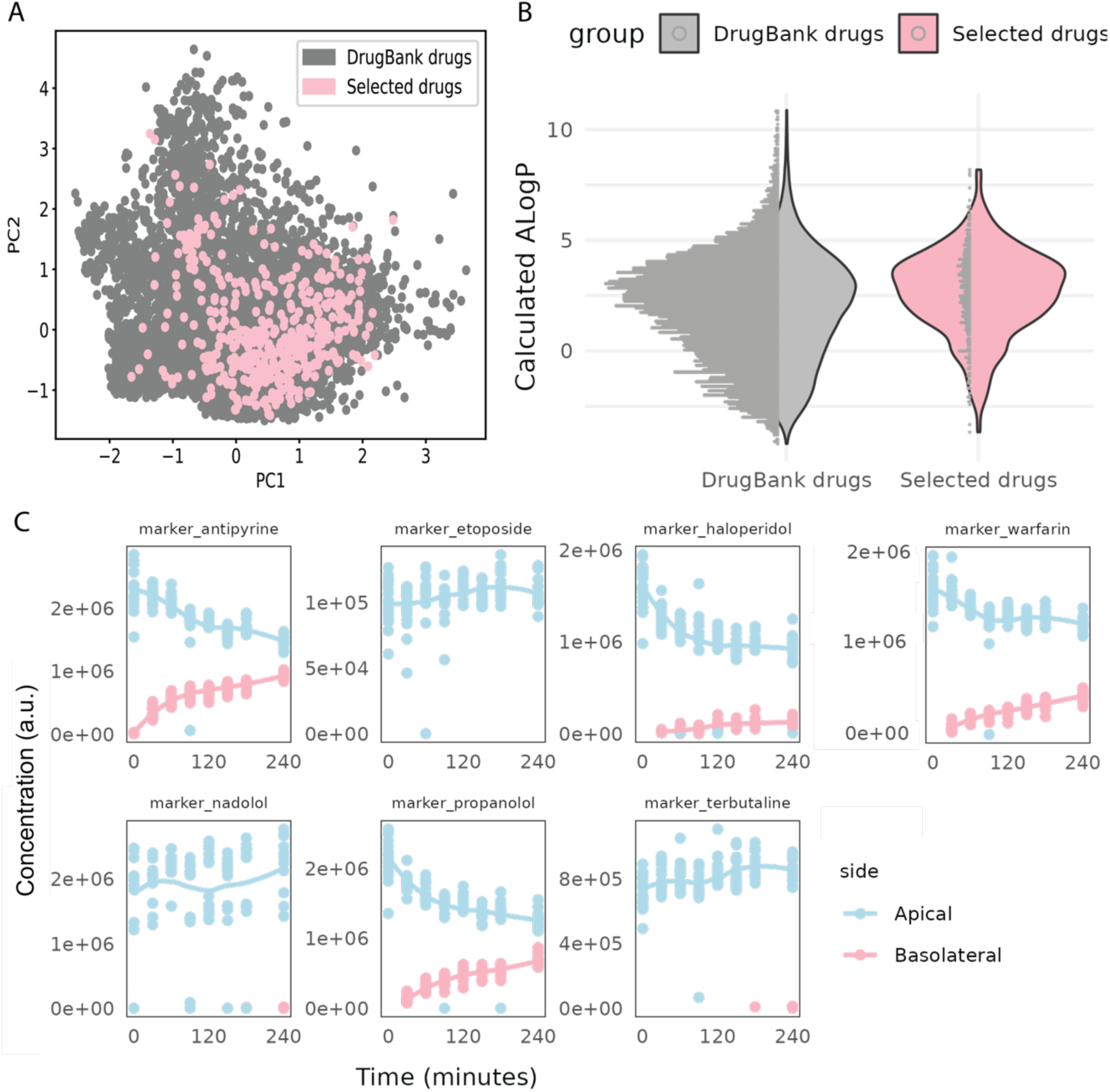
A. Chemical diversity of the 482 selected drugs (red dots) compared to 11583 drugs from DrugBank (grey dots). B. Comparison of distribution of calculated LogP (CLogP) values between drugs from DrugBank and the drugs selected in our test panel. C. Marker compound kinetics in the apical and basolateral compartment of transwells with Caco2 monolayers exposed to sterile bacterial cuture media on the basolateral side (n=4).

**Supplementary figure 4:**
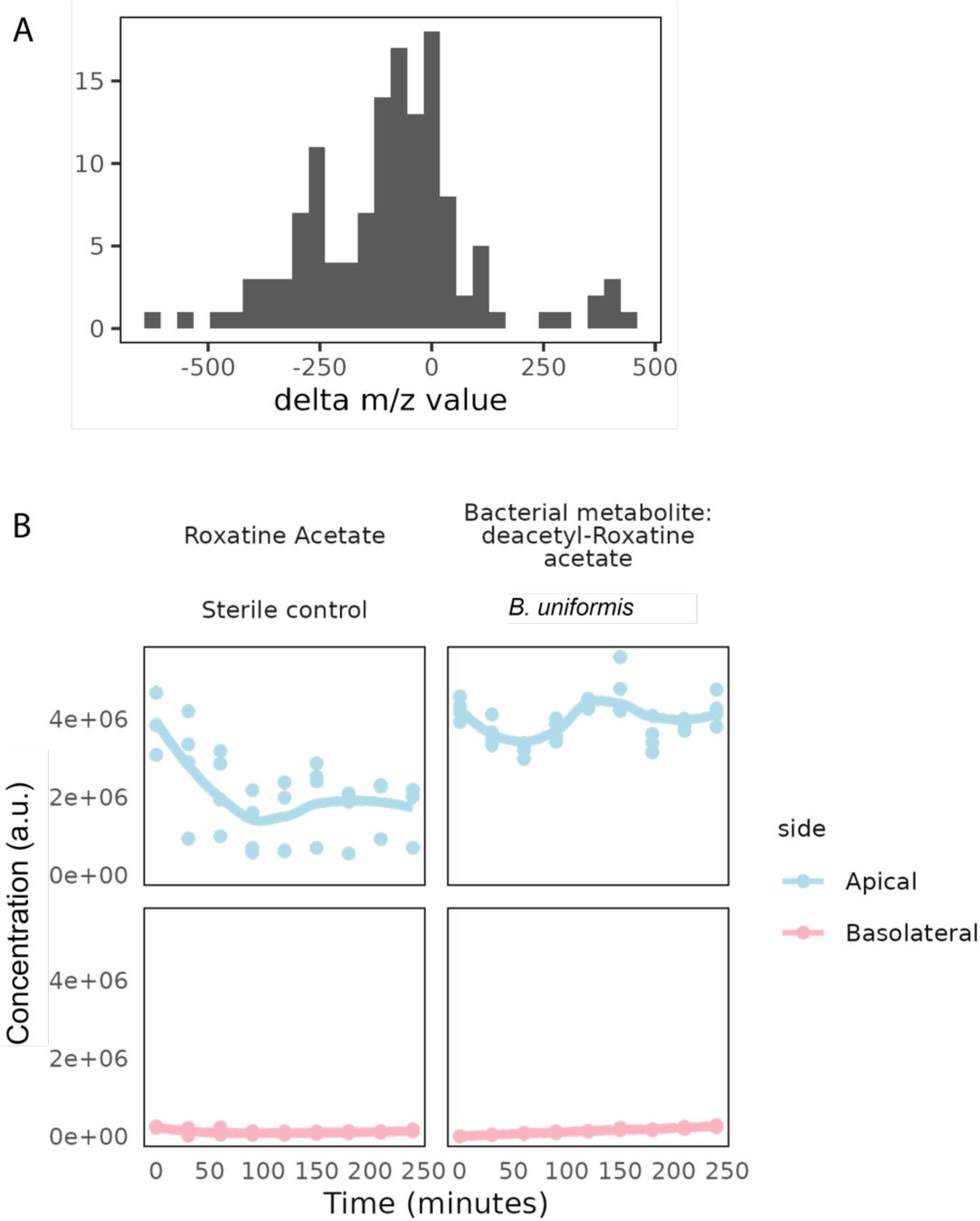
A. Histogram of mass shifts between the parent drug and the putative drug metabolites. B. Permeability of roxatidine acetate and its known bacterial metabolite, roxatidine, in the transwell assay (n = 4).

**Supplementary figure 5:**
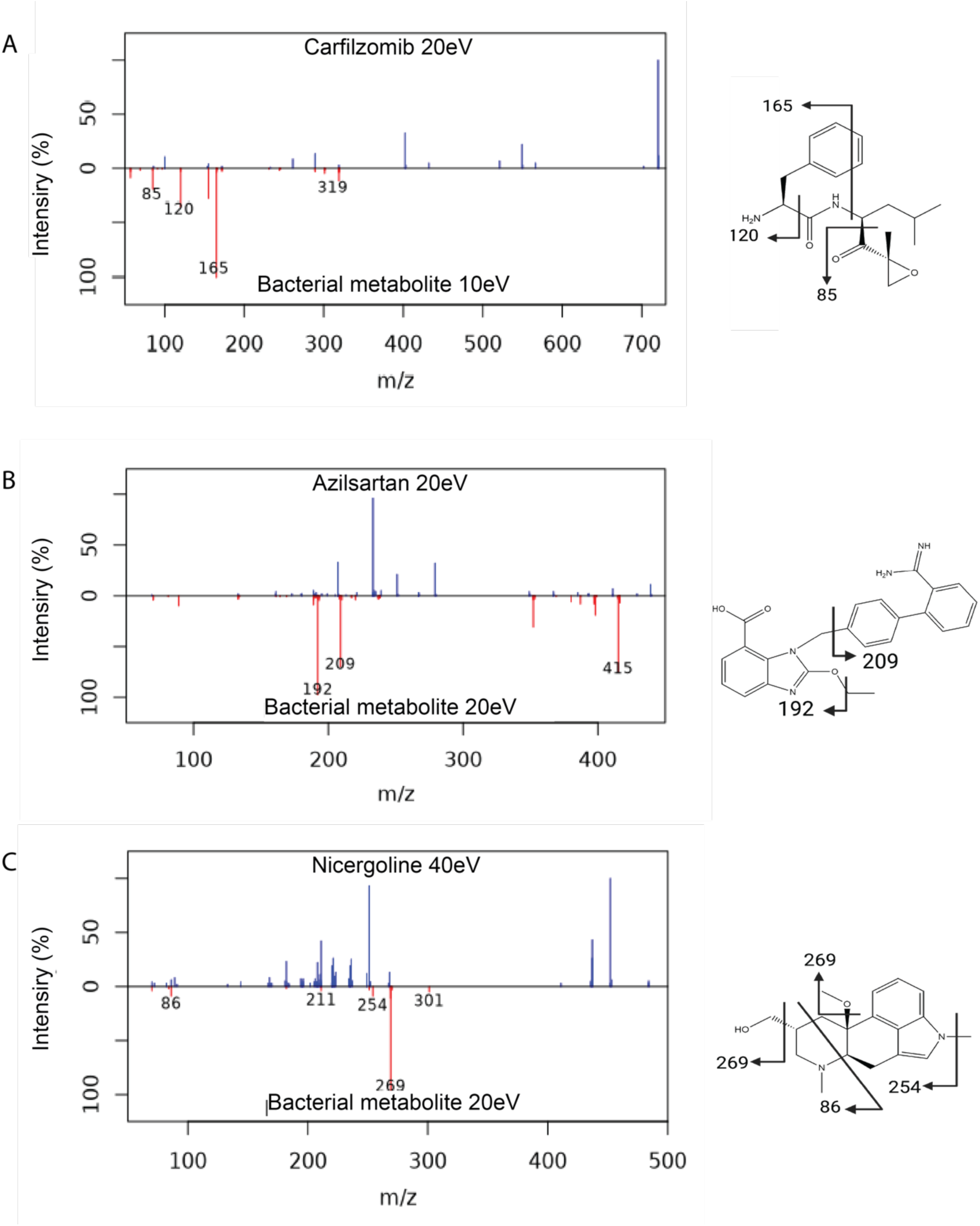
A. MS/MS spectra of the drug carfilzomib and its bacterial metabolite. B. MS/MS spectra of azilsartan and its bacterial metabolite. C. MS/MS spectra of nicergoline and its bacterial metabolite

